# FORAGING MECHANISMS IN EXCAVATE FLAGELLATES SHED NEW LIGHT ON THE FUNCTIONAL ECOLOGY OF EARLY EUKARYOTES

**DOI:** 10.1101/2023.11.20.567814

**Authors:** Sei Suzuki, Federica Miano, Seyed Saeed Asadzadeh, Alastair G.B. Simpson, Thomas Kiørboe

**Author notes:** Equal authorship (Sei Suzuki, Federica Miano).

## Abstract

The phagotrophic flagellates described as ‘typical excavates’ have been hypothesized to be morphologically similar to the Last Eukaryotic Common Ancestor and understanding the functional ecology of excavates may therefore help shed light on the ecology of these early eukaryotes. Typical excavates are characterized by a posterior flagellum equipped with a vane that beats in a ventral groove. Here, we combined flow visualization and observations of prey capture in representatives of the three clades of excavates with computational fluid dynamic modelling, to understand the functional significance of this cell architecture. We record substantial differences amongst species in the orientation of the vane and the beat plane of the posterior flagellum. Clearance rate magnitudes estimated from flow visualization and modelling are like that of other similarly sized phagotrophic flagellates. The interaction between a vaned flagellum beating in a confinement is modelled to produce a very efficient feeding current at low energy costs, irrespective of the beat plane and vane orientation and of all other morphological variations. Given this predicted uniformity of function, we suggest that the foraging systems of typical excavates studied here may be good proxies to understand those potentially used by our distant ancestors more than 1 billion years ago.

**Significance:** Human sperm reminds us of our ancestry: flagellates, unicellular organisms equipped with a flagellum. The last common eukaryotic ancestor (LECA) was a flagellate. Phylogenetic analyses suggest that Excavates, an assemblage of flagellates, are the living organisms most similar to LECA. They have distinct characteristics in common: a ventral groove within which a vaned flagellum is beating. We show how the shared morphology and foraging behavior among 3 excavate clades is fluid dynamically efficient. A similar flagellar arrangement, potentially homologous to that found in the excavates, is found among flagellates from other deep branches of the eukaryotic tree, suggesting that the typical excavate foraging system studied here may have been used by our distant ancestors more than 1 billion years ago.

---

Phagotrophic flagellates are the main consumers of bacteria and pico-phytoplankton in the ocean, and thus play a key role in regulating pelagic microbial ecosystems (1). They are phylogenetically very diverse, with representatives in all the major branches of the eukaryotic tree of life, and thus central to the evolution of all eukaryotes (2). They are also functionally diverse, as the number of flagella, their wave patterns and kinematics, and whether their flagella are naked or equipped with hairs or vanes vary between groups (3). Many species can also photosynthesize, i.e., they are mixotrophic. This functional diversity may imply great variation in foraging efficiency, as quantified by clearance rates that vary by more than 2 orders of magnitude between flagellates of similar sizes (4).

The mechanisms of phagotrophic foraging have been examined in only a few groups of ecologically important flagellates. In the small-scale world of flagellates where the Reynolds number (ratio of inertial to viscous forces) is low, viscosity impedes predator-prey contact, and generating a sufficient feeding current typically requires forces that are larger than what a single naked flagellum can produce (5). Thus, in the groups examined in detail, one finds that the flagellar apparatus has highly specialized adaptations for foraging. For example, Stramenopiles have one flagellum equipped with rigid hairs that increases the force production of the beating flagellum by a factor of 5-10 (6, 7), and choanoflagellates have their single flagellum equipped with a vane that allows it to pump water through their collar filter (8, 9). Dinoflagellates are equipped with a transverse flagellum embedded in a ‘sock’ and runs in a groove around the cell circumference (10), which allows a very powerful feeding current (5). These are all groups that are quantitatively important in the water column.

A less appreciated (and phylogenetically less coherent) assemblage of phagotrophic flagellates that are important for understanding deep eukaryote evolution are the ‘typical excavates’. They occur in the water column mainly attached to marine snow particles as well as in sediments as well (11–16), and they may be abundant in some anoxic marine water masses (16, 17). Like the phagotrophic flagellates mentioned above, the typical excavates also have a specialized flagellar apparatus. The best-known forms are biflagellated cells with a naked anterior flagellum, and a posterior flagellum that beats in a ventral groove and bears 1-3 broad vanes of unclear function (16, 18) (Fig. 1a). Each vane is formed by an intraflagellar (paraxonemal) protein lamella that distends the flagellar membrane, quite unlike the fine extracellular filamentous vanes of choanoflagellates mentioned above. Bacterial prey are collected and engulfed in the posterior end of the groove, but the prey concentration- and foraging mechanisms are essentially unknown.

**Fig. 1.**
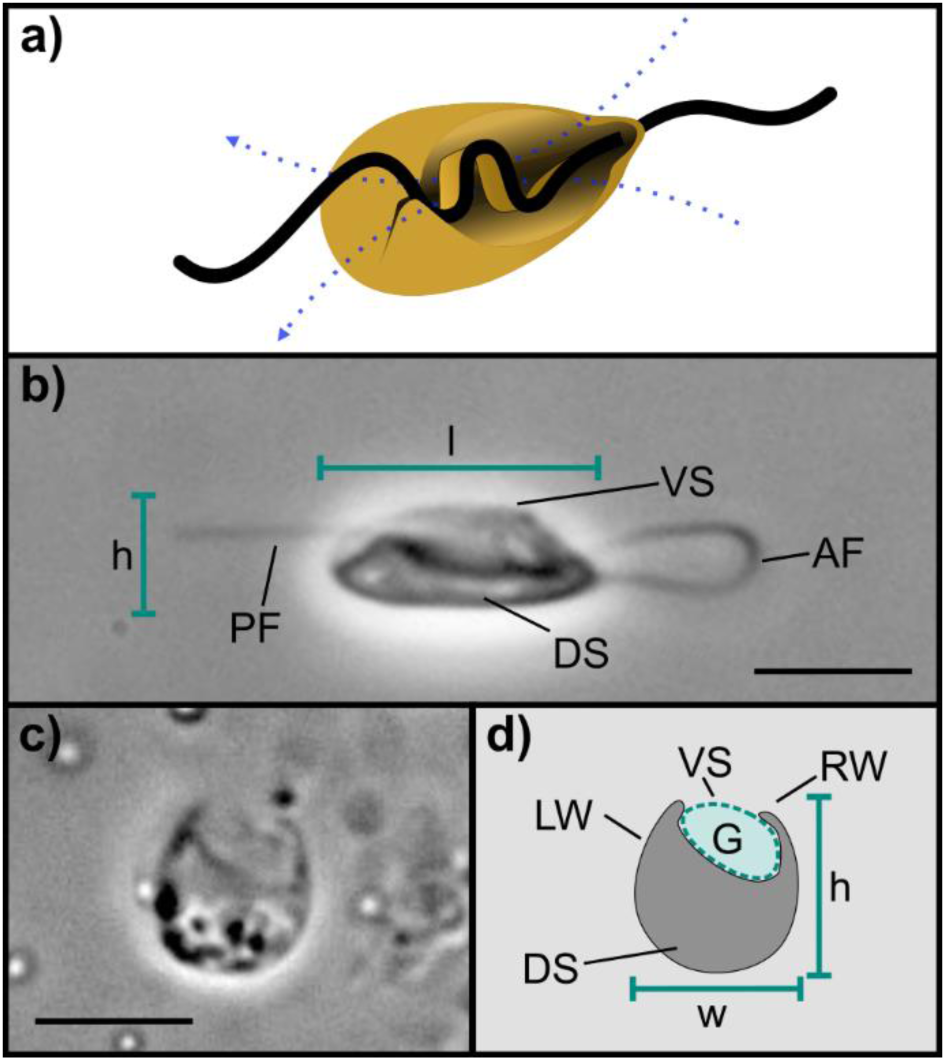
‘Typical excavate’ flagellate and morphometric parameters: a) ‘typical excavate’ flagellate with an anterior flagellum, and a posterior flagellum with a vane that runs in the ventral groove (dashed blue arrows indicate the feeding flow lines); b) Live image of the left longitudinal side of the cell (*Jakoba libera*); c) Live image of the ventral groove focused at mid cell body length (*Jakoba libera*, edited image: horizontal flip); d) schematic drawing of image c. Abbreviations: h = cell height, l = cell length, w = cell width, PF = posterior flagellum, AF = anterior flagellum, VS = ventral side, DS = dorsal side, RW = right-side groove wall, LW = left-side groove wall, and G = groove area (marked with dashed line). Scale = 5 μm.

In addition to their ecological role in the modern ocean, excavates are of great evolutionary interest because of the possibility that the Last Eukaryotic Common Ancestor (LECA) resembled living typical excavates (19). This is a highly consequential hypothesis for understanding the evolutionary origins and history of modern complex cells. Understanding the mechanisms of swimming and foraging in the excavate flagellates may therefore help us infer how LECA cells used their flagella to feed and swim in the ancient past.

Here we examine the mechanisms and fluid dynamics of foraging in several species of typical excavates belonging to the three major groups: Discoba, Metamonada, and Malawimonadida. We use high speed video-microscopy to describe the beating of the flagella and capture of prey, particle tracking to visualize the feeding current and to estimate clearance rates, and computational fluid dynamics to understand the functional significance of the characteristic morphological features of excavate cells.

## Material and Methods

### Culturing

The studied flagellates were laboratory cultures maintained in the Simpson Lab, Dalhousie University (*Carpediemonas membranifera* isolate BICM), or kindly provided by Franz Lang, Université de Montréal (*Reclinomonas americana* isolate ATCC50394, *Jakoba libera* isolate ATCC50422, ‘*Malawimonas californiensis*’ isolate ATCC50740) or Julie Boisard and Courtney Stairs, University of Lund (*Kipferlia bialata* isolate WC1A). Cultures were maintained in the dark at 18°C in highly diluted Miller’s LB broth media to feed the naturally occurring bacteria that served as prey. The aerobic freshwater species, *R. americana*, grew with 0.3% LB in Milli-Q® water; the aerobic marine species J. *libera* and *M. californiensis* grew with 0.3% LB in filtered and pasteurized North Sea water (salinity 30‰); and the anaerobic marine species *C. membranifera* and *K. bialata* grew in 3% LB in sterilized North Sea water under near-anoxia established by the prokaryotic growth on this richer medium.

### Microscope imaging

Phase contrast imaging was performed with an Olympus IX71 inverted microscope equipped with an Olympus UPLanFL N oil immersion x100 / 1.30 objective and an attached high-speed *Phantom Camera* (Miro LAB 320). Videos had a minimum resolution of 512 x 512 pixels and were recorded at 300 or 500 frames per second (fps). The effects of light heating were mitigated with short exposure intervals (< 10 minutes) at moderate intensities during recordings. Videos and images were later analysed using the open-source software ImageJ (Fiji).

Aerobic flagellate species were observed by mounting 100 µL of culture between two glass coverslips (24 x 24 mm) that were spaced by small blobs of Vaseline® in each corner. The observation chamber for anaerobic species consisted of a plastic ring (16 mm inner diameter and 1.5 mm height) fixed with Vaseline® to a coverslip, and filled with 300 µL of culture, then sealed on top with a second coverslip, held by surface tension. The latter ring chamber set-up was also used for all species when visualising their flow fields with tracer particles (see below).

### Morphometrics

Morphometrics and cross-sectional areas of the groove were estimated from live images of feeding cells extracted from videos (Fig. 1b, c, d), assisted by TEM images from the literature (11, 13, 20) (Table 1). Volumes of individual flagellate cells were estimated using a Coulter Counter.

**Table 1.** Cell body and ventral groove morphometrics.

|  | Cell body |  |  |  |  |  |  |  |  | Groove |  |  |  |  |  |  |  |
| --- | --- | --- | --- | --- | --- | --- | --- | --- | --- | --- | --- | --- | --- | --- | --- | --- | --- |
|  | Length (μm) |  | Height (μm) |  | Width (μm) |  | Depth (μm<sup>2</sup>) |  | Volume (μm<sup>3</sup>) | Length (μm) |  | Width (μm) |  | Depth (μm) |  | Area (μm<sup>2</sup>) |  |
|  | Mean ± SD | n | Mean ± SD | n | Mean ± SD | n | Mean ± SD | n |  | Mean ± SD | n | Mean ± SD | n | Mean ± SD | n | Mean ± SD | n |
| *Jakoba libera* | 7.8 ± 0.6 | 7 | 3.7 ± 1.0 | 6 | 4.4 ± 0.7 | 3 | 3.3 ± 0.0 | 1 | 24 | 6.3 ± 0.0 | 1 | 2.9 ± 0.3 | 3 | 2.1 ± 0.0 | 1 | 5.0 ± 0.0 | 1 |
| *Reclinomonas americana* | 7.8 ± 0.7 | 6 | 3.5 ± 0.2 | 2 | 5.5 ± 0.9 | 5 | 4.2 ± 0.5 | 3 | 58 | 5.3 ± 0.0 | 1 | 3.7 ± 0.5 | 3 | 2.8 ± 0.5 | 3 | 9.6 ± 1.7 | 3 |
| *Kipferlia bialata* | 9.3 ± 0.6 | 5 | 5.4 ± 0.5 | 3 | 5.2 ± 0.0 | 1 | 2.9 ± 0.9 | 4 | 102 |  |  | 3.3 ± 0.0 | 1 | 1.8 ± 0.0 | 2 | 5.6 ± 1.1 | 2 |
| *Carpediemonas membranifera* | 7.2 ± 0.7 | 7 | 4.4 ± 0.7 | 5 |  |  | 2.0 ± 0.0 | 2 | 28 | 5.3 ± 0.0 | 1 | 2.9 ± 0.4 | 6 | 2.7 ± 0.2 | 2 | 3.8 ± 0.0 | 2 |
| *Malawimonas californiensis* | 5.4 ± 0.4 | 6 | 3.1 ± 0.5 | 3 | 3.1 ± 0.5 | 3 | 2.0 ± 0.0 | 1 |  |  |  | 2.3 ± 0.1 | 2 | 1.4 ± 0.0 | 1 | 2.3 ± 0.0 | 1 |

### Flow fields

The ring chamber was filled with a dilute suspension of flagellates, and a single cell was focused under the microscope. Then, 0.5-µm tracer particles were gently added to the chamber sealed with a coverslip. Videos of each of four species (*J*. *libera*, *R*. *americana*, *K. bialata*, and *C*. *membranifera*) were selected for analyses and particles were tracked at a frequency of 50 or 60 Hz using the Manual Tracking function in ImageJ. (e) Clearance rates Clearance rates were estimated on the assumption that particles transported into the ventral groove can be considered captured, and hence the clearance rate can be estimated as the volume flow rate in the groove. We estimate flow velocities in the groove by noting the time and position of particles as they entered the groove and either left it or stopped at its rear end. Here, most particles were eventually discarded, likely because the flagellate could not phagocytize further particles due to a full food vacuole. The clearance rate was then estimated as the flow velocity multiplied by the cross-sectional area of the groove.

### Computational Fluid dynamics

Computational fluid dynamics (CFD) modelling allows one to quantify the flow field generated by a beating flagellum, given known morphology, wave patterns, and beat kinematics, by numerically solving the flow-governing Navier-Stokes equations. We developed a CFD model of a simplified generic excavate flagellate (Fig. S1) with basic morphology and kinematics inspired by *Jakoba libera* (Tables 1, 2); that is, a cell with a posterior flagellum beating inside and parallel to the bottom of the ventral groove and equipped with an inward oriented vane (See online material for details). This basic case was then modified to have other vane arrangements (outward oriented, broader vane, two vanes), beat orientation (perpendicular to the bottom of the groove), a higher wall on the right than on the left side of the groove (Fig. S2), attachment to a surface positioned on the dorsal or ventral side of the cell, an anterior flagellum beating in a 3-dimensional pattern, and/or a posterior flagellum that is also active outside the ventral groove, as found in the different species described below. We varied one parameter at the time to examine the functional significance of the different morphological features found in the different species. Finally, to examine the effect of having the posterior flagellum constrained in a groove, we ran a case without any groove and the flagellum instead beating above the surface of the cell (Fig. S3). As diagnostic output characteristics we computed flow fields, power consumption, and magnitudes of the clearance rate.

**Table 2.** Beat frequencies of flagella in feeding (attached) and free-swimming flagellates and swimming speeds.

|  | Feeding |  |  |  | Free-swimming |  |  |  |  |  |
| --- | --- | --- | --- | --- | --- | --- | --- | --- | --- | --- |
| | Posterior Flagellum<br>beat frequency (Hz) | | Anterior Flagellum<br>beat frequency (Hz) | | Posterior Flagellum<br>beat frequency (Hz) | | Anterior Flagellum<br>beat frequency (Hz) | | Speed ($\mu\text{m s}^{-1}$) | |
| | Mean ± SD | n | Mean ± SD | n | Mean ± SD | n | Mean ± SD | n | Mean ± SD | n |
| *Jakoba libera* | 37 ± 3 | 6 | - | - | 32 ± 0.5 | 2 | 17 ± 2 | 2 | 15 ± 3 | 2 |
| *Reclinomonas americana* | 49 ± 4 | 5 | 48 ± 4 | 5 | 60 ± 5 | 10 | 44 ± 3 | 11 | 105 ± 12 | 8 |
| *Kipferlia bialata* | 26 ± 3 | 4 | 6 ± 2 | 5 | 24 ± 19 | 2 | 10 ± 2 | 2 | 12 ± 5 | 2 |
| *Carpediemonas membranifera* | 27 ± 2 | 5 | 6 ± 0 | 5 | 31 ± 4 | 2 | 7 ± 1 | 2 | 27 ± 13 | 2 |
| *Malawimonas californiensis* | 42 ± 6 | 5 | 7 ± 1 | 5 | | | | | | |

The Navier-Stokes equations were discretized using a finite volume method and solved on a discrete representation of the computational domain composed of polyhedral cells (Fig. S4). We utilized the commercial CFD software STAR-CCM+ (version 18.02.008-R8) for the numerical simulations. We used mesh morphing along with the overset method to move the computational mesh for flagellar motion. The morphing method deforms the computational mesh around the flagella in response to their prescribed motion. With the overset method, rather than moving the entire computational mesh, it deforms the mesh only around the flagella. A stationary background mesh is overlapped with the deforming mesh, and field data is interpolated between them for a smooth solution (see online supplementary material for a full technical description).

## Results

### Feeding behaviour and free-swimming

Prey capture and handling follows a general pattern shared among all the species examined. The ventral groove extends for most of the length of the cell, with its right wall generally being taller and more continuous. A feeding current is generated by the flagella, always involving the posterior vane-bearing flagellum beating in the groove region with high amplitude waves at 25-50 Hz (Table 2). The behaviour of the anterior flagellum differs among species.

The flow drives the prey into and through the ventral groove, entering roughly in the middle, and draws it towards the posterior end of the cell. Food particles are captured in the groove and handled near the phagocytosis site at its posterior end. If the prey is rejected, it will exit the groove. The posterior flagellum sometimes actively retains the particle by pushing it against the right wall of the groove when it stops beating. Meanwhile, a moving ‘wave’ of the cell membrane is observed in *K. bialata* and *R. americana* – often as a transversal slender dark shadow - sweeping posteriorly along the groove and disappearing when the wave reaches the phagocytosis site in the posterior end of the groove (Movie S2 and S3). When the wave reaches the end of the groove, the particle is ingested with the membrane observed to affect the closure of the phagocytic vacuole.

When feeding, *Jakoba libera* attaches to a surface with a bent anterior (Fig. 2a). Often, the anterior flagellum will flex repeatedly and push the body backwards and forwards or change its orientation (Movie S1). The feeding current is generated by the posterior flagellum only. It extends a third of a body length outside of the cell and this portion is straight and rigid. When swimming, the anterior flagellum is mainly responsible for propulsion as it beats in a three-dimensional power-and-return stroke while the posterior flagellum remains beating inside the groove (Movie S1).

**Fig. 2.**
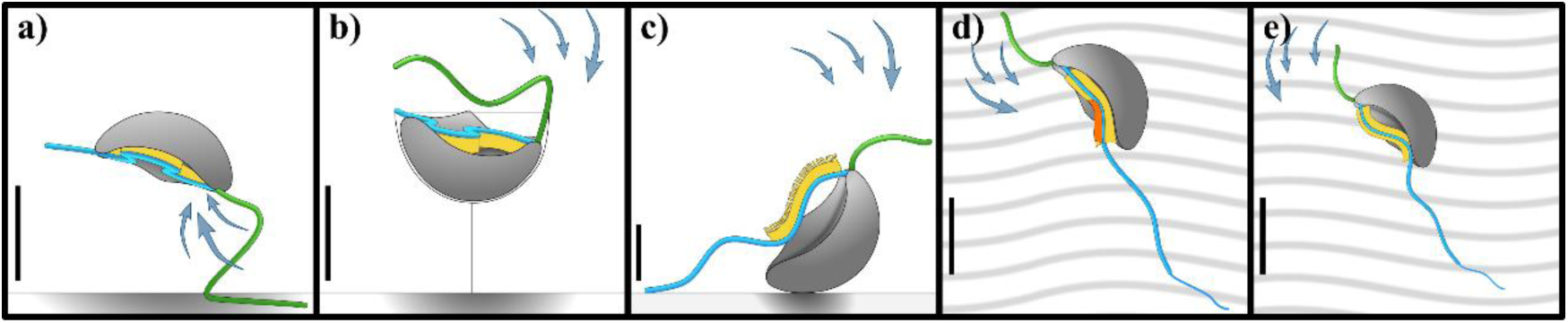
Schematic representations of typical excavate species: a) *Jakoba libera*, b) *Reclinomonas americana*, c) *Kipferlia bialata*, d) *Carpediemonas membranifera*, and e) *Malawimonas californiensis*. While foraging, *J. libera, R. americana,* and *K. bialata* are attached to the surface (bottom line); instead, *C. membranifera* and *M. californiensis* skid on the substrate (wavy background). Each species panel includes cell body (gray), anterior flagellum (blue), posterior flagellum (green), feeding current direction (arrows), and flagellar vane number and position (yellow). Note that *C. membranifera* has a short third vane (orange). Reference scale bar = 5 µm.

In its feeding stage, *Reclinomonas americana* sits inside a lorica that is attached to the substrate with the ventral groove facing up (Movie S2) (Fig. 2b). It has a very tall and pronounced right wall. Uniquely, both flagella beat rapidly (Table 2). The anterior flagellum beats in a three-dimensional pattern above the ventral groove with the distal end pointing posteriorly. A short, straight portion of the posterior flagellum extends posteriorly beyond the groove and moves rigidly.

After cell division, one daughter cell remains in the parental lorica, while the other is free-swimming (12, 21). This swarmer is slender and highly mobile (Tables 1 and 2) (Movie S2). Its anterior flagellum performs broader power-and-return strokes in front of the cell with the distal tip always pointing posteriorly. The posterior flagellum is longer (approximately 2 body lengths) and trails freely, beating along the groove.

*Kipferlia bialata* attaches to a surface with the posterior end of the cell body, normally with the groove facing away from the surface and generating a feeding current with both flagella (Movie S3) (fig. 2c). The anterior flagellum performs slow (<10 Hz) three-dimensional power-and-return strokes. The posterior flagellum also beats in in 3 dimensions but beats much faster (∼30 Hz). When free-swimming, the flagellar kinematics remain the same. The anterior flagellum is responsible for displacement and causes the cell body to rotate on its longitudinal axis (Movie S3).

*Carpediemonas membranifera* and *Malawimonas californiensis* have similar predation behavior and flagellar kinematics (Movie S4) (Fig.s 2d, 2e). When feeding, the cell attaches to surfaces with the distal ∼quarter of the posterior flagellum and the cell hovers or skids around while generating a feeding current with both flagella. The anterior flagellum executes slow power-and-return strokes similar to *K. bialata*. The posterior flagellum beats only with the portion that is inside the groove. *C. membranifera* does not rely solely on feeding currents to capture prey, being recorded as ‘scooping’ inside the groove a particle that was attached to the surface through movement of the cell body (Movie S4). Swimming cells use the posterior flagellum for propulsion (Movie S4). The overall contribution of the anterior flagellum to motility is minor and possibly more dedicated steering.

### The ventral groove and the flagellar vane

The portion of the posterior flagellum inside the groove is equipped with vanes making much of the flagellum itself paddle-shaped. These thin extensions of the flagellum double, at least, its surface area and are somewhat flexible. We observed the number of vanes, their orientation, and their spatial-temporal distribution at the mid cross section of the groove (Movie S5) (Fig. 3). Three functional types were observed: 1) single inward vane, moving parallel to the groove floor, 2) single outward vane, moving in a 3D motion, and 3) double vanes. Irrespective, the vane(s) project more-or-less perpendicularly to the primary plane of beating.

1. Single inward vane (*Jakoba libera* and *Reclinomonas americana*) The posterior flagellum of *J. libera* and *R. americana* remains inside the groove throughout the beat cycle. It beats in an oblique plane between the two walls: from the top of the left-side wall towards the bottom of the right-side wall (Movie S5) (Fig. 3a). The taller right wall curves over the groove. The flagellar beating plane is parallel to the floor and the is vane sweeping across it in proximity or in direct contact.
2. Single outward vane (*<u>Kipferlia bialata</u>*) The flagellum has a terminal row of tape-like hairs (22). The beating pattern is three-dimensional, spanning from the bottom to outside of the groove (Movie S5) (Fig. 3b). The initial left-side tilt of the vane becomes almost parallel to the groove floor when it collides with the base of the left wall, and it subsequently points outwards when it emerges outside of the groove again at the end of the beat cycle.
3. Double vanes (*Carpediemonas membranifera* and *Malawimonas californiensis*) The outwards vane has a slight tilt to the left while, symmetrically, the inwards vane points to the right. The beat wave oscillates from just outside of the groove to the bottom-left corner of the cavity, executing an oblique trajectory between the walls (Movie S5) (Fig. 3c). A third, much shorter vane, orthogonal to the two main vanes, has been reported for *C. membranifera* by electron microscopy (13).

**Fig. 3.**
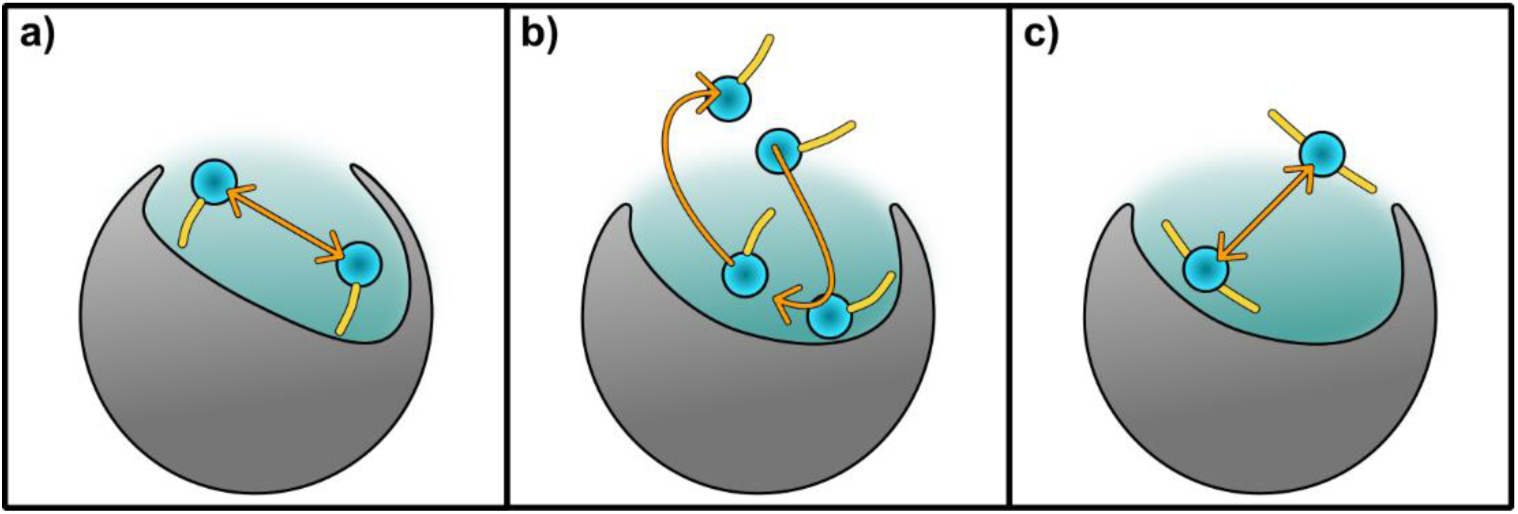
Posterior flagellum kinematics inside the ventral groove. Schematic representations of mid-transversal cross section views of the ventral groove (teal shadow) and the position of the flagellum (blue circle) equipped with a vane/s (yellow) between the maximum beat wave amplitudes (arrows). The right-side wall of the cell is on the right-side of the viewer. The three cross sections correspond to the observations of: a) *Jakoba libera* and *Reclinomonas americana*; b) *Kipferlia bialata*; c) *Carpediemonas membranifera* and *Malawimonas californiensis*.

### Feeding current and clearance estimates

Example particle tracks for 4 species represent two-dimensional projections of three-dimensional flows and depend also on what angle the cell was observed (Fig. 4). Thus, the particle tracks can be used only to qualitatively describe the feeding current. The feeding current flows are similar among all species, with an accelerating flow arriving at the mid-to-anterior end of the ventral groove and leaving again from the mid-to-posterior end of the groove. Far from the ventral groove, particle motion is dominated by Brownian diffusion, and the feeding current only dominates particle motion within 10-30 µm of the cell.

**Fig. 4.**
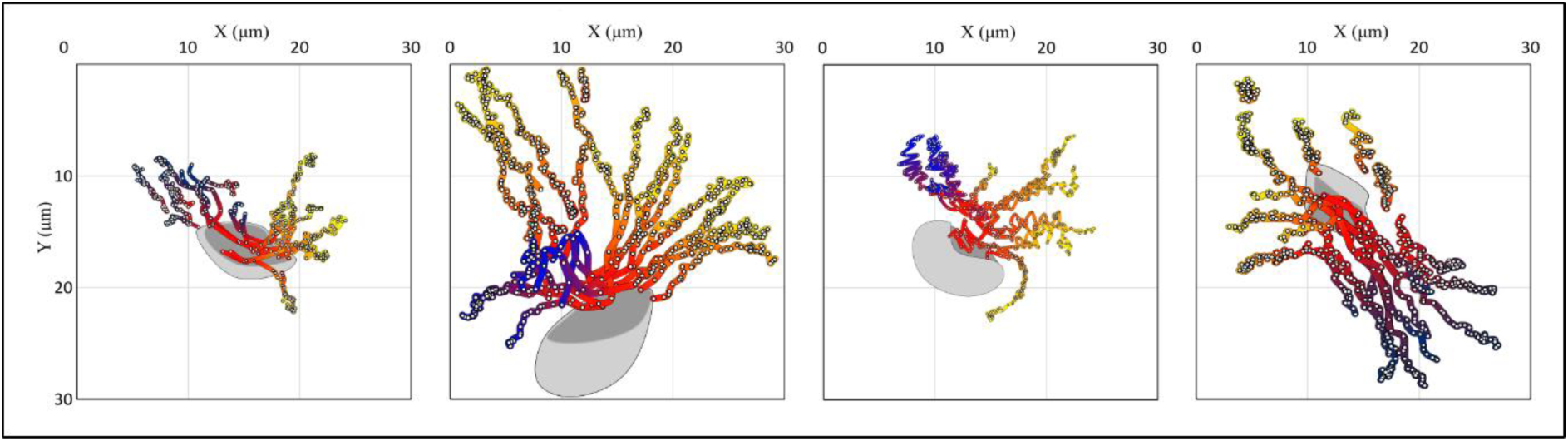
Example particle tracks for 4 species: a) *Jakoba libera*, b) *Reclinomonas americana,* c) *K. bialata,* and d) *C. membranifera,*. Tracer particles are positioned (dots) at a frequency of 60 Hz (Except R. *americana*: 50 Hz), and the distance between positions thus indicates flow speed. Far from the flagellate particle tracks are dominated by Brownian motion. The color indicates the direction of the flow, from yellow through red to blue.

Flow velocities in the ventral groove varied between species by a factor of ca. 3, apparently unrelated to the beat frequency of the posterior flagellum (Table 2). The resulting estimates of cell-volume specific clearance rates vary by an order of magnitude, between 0.2-3 x 10^6^ per day (Table 3).

**Table 3.**
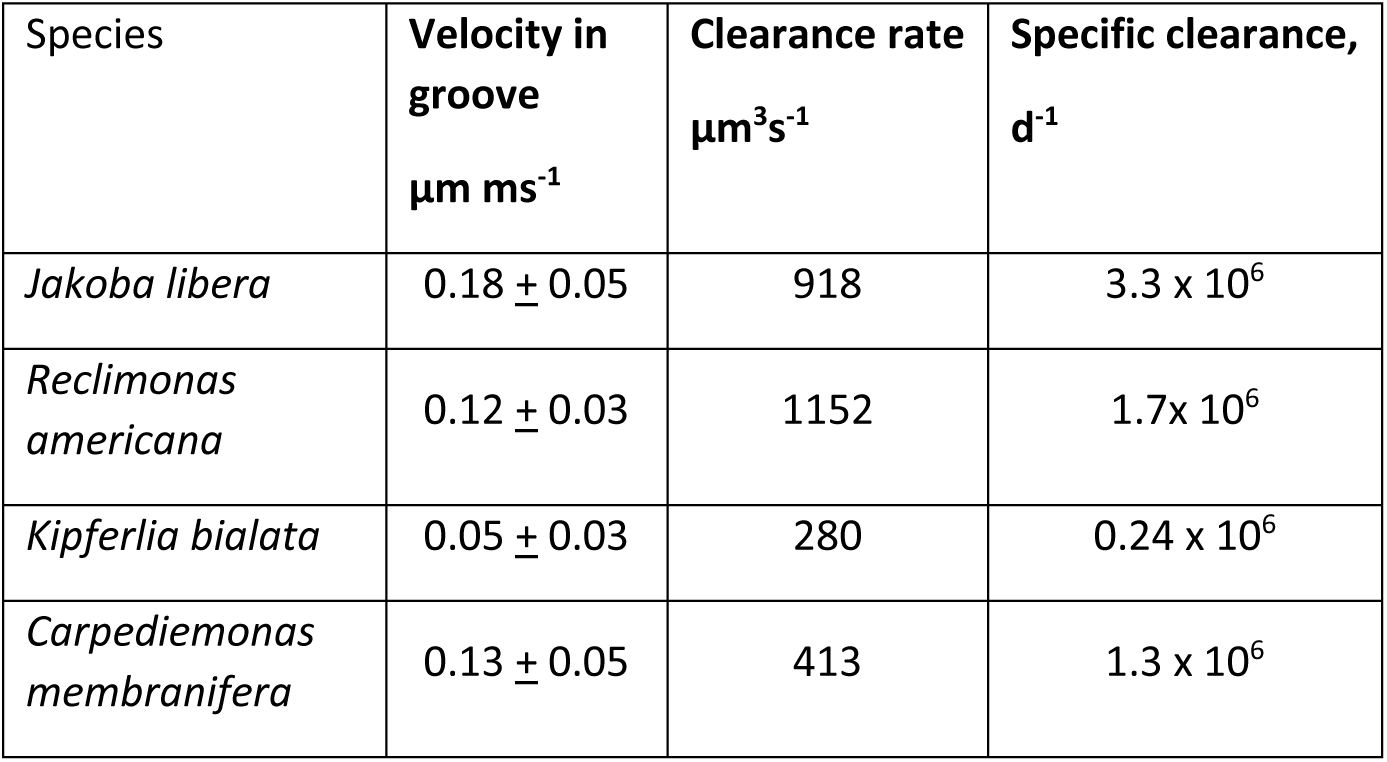
Estimated clearance rates. Specific clearance rate is the clearance rate normalized by cell volume.

### CFD

The main outcomes of the CFD modelling are first that the magnitude of the clearance rate depends mainly on the beating of the posterior flagellum inside/over the ventral groove and on the dimensions of the vane, and that all other morphological features as well as the action of the anterior flagellum have only limited effect (Fig. 5). The model flagellates have clearance rates of 250 - 450 μm^3^s^-1^, of similar order to that estimated experimentally for the examined species (Fig. 5). Whether the posterior flagellum beats parallel or perpendicular to the bottom of the groove has limited effect, so both orientations observed appear equally well suited for feeding. Broader vanes lead to higher power consumption, but also to higher clearance rates.

**Fig. 5.**
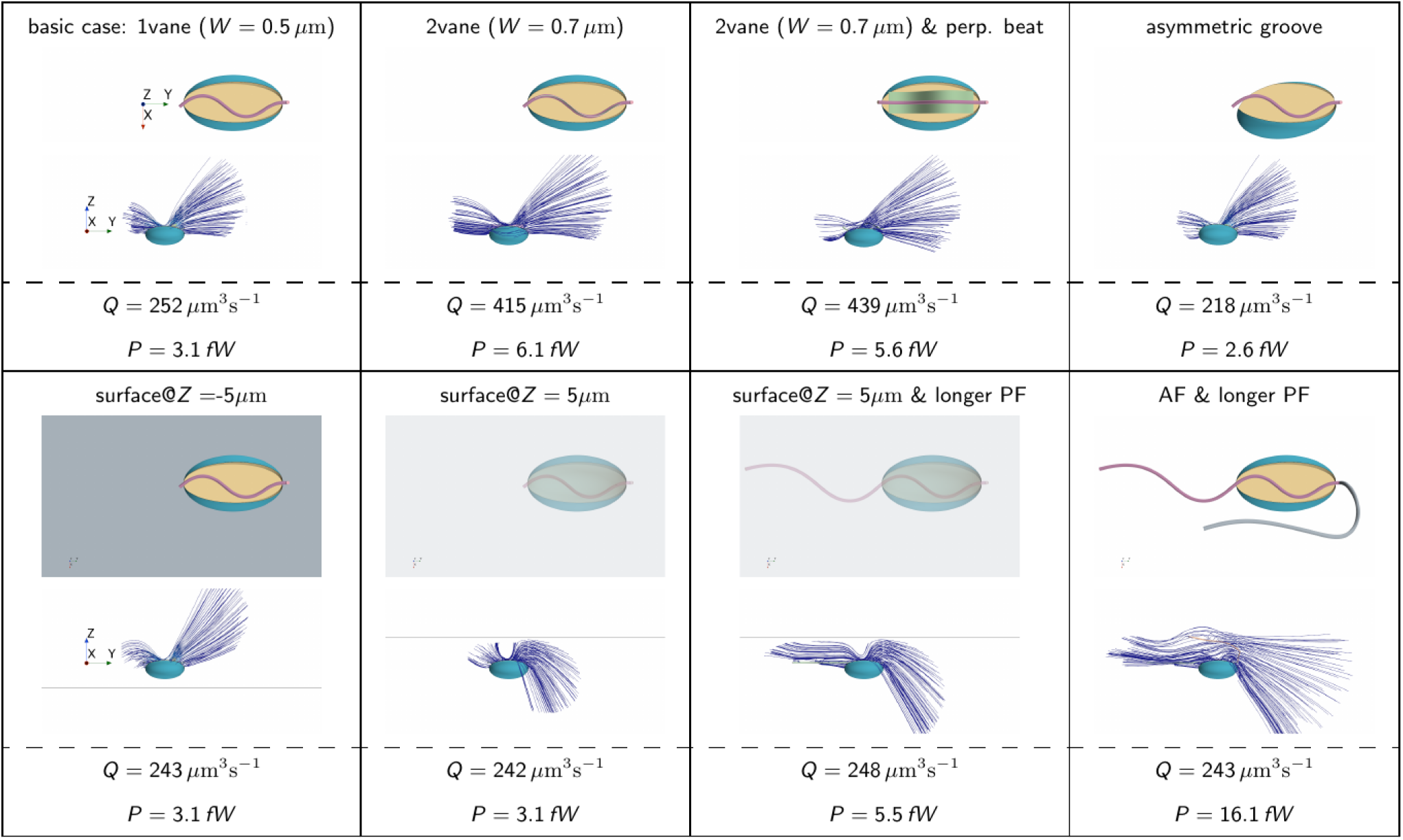
CFD cases demonstrating the influence of various morphological aspects and surface proximity. In each panel the case is first described (number of vanes, width of vane, asymmetry of groove, the presence of a surface dorsally (−5 µm) or ventrally (+5 µm) to the cell, and the presence of an active anterior and extended posterior flagellum. The anterior end of the cell is to the right. Streamlines represent the averaged flow field, with sections omitted where flow velocity is below 2 µm s^-1^ (corresponding to a Peclet number of ∼1 for 0.5 µm passive prey particles). This threshold indicates where the advective feeding current overcomes the diffusive Brownian motion of passive prey particles. *Q* is the estimated clearance rate and *P* the estimated power consumption.

Second, the presence of the anterior flagellum and the ‘free’ part of the posterior flagellum extending outside the groove improves the architecture of feeding current by extending the region around the cell within which the advective feeding current is strong enough to overcome the diffusive Brownian motion of passive prey particles. This is illustrated by longer streamlines that are plotted only where the Peclet number (Pe) exceeds 1 (Fig. 5); The Peclet number indicates the relative significance of advection over diffusion, Pe = *au*/*D*, where *a* is prey radius (0.5 µm), *D* is Brownian diffusion computed from Einstein relation, and *u* is the feeding current velocity). This enhancement of the feeding current is also obvious in our observations (Fig. 4) and is particularly significant for flagellates attached to a surface (Fig. 5 5). The improvement, however, happens at the cost of higher power consumption.

Finally, the containment of the vaned flagellum in the ventral groove improves the magnitude of clearance rate. A vaned flagellum beating outside a groove creates a feeding current characterized by intense sideways oscillations that cancel out during a complete beat cycle, resulting in a lower (averaged) clearance rate. Conversely, the presence of a groove generates a smoother and more directional flow, leading to a higher clearance rate (Fig. 6). The advantage of beating inside a groove vanishes for a naked flagellum without a vane (Fig. 6c).

**Fig. 6.**
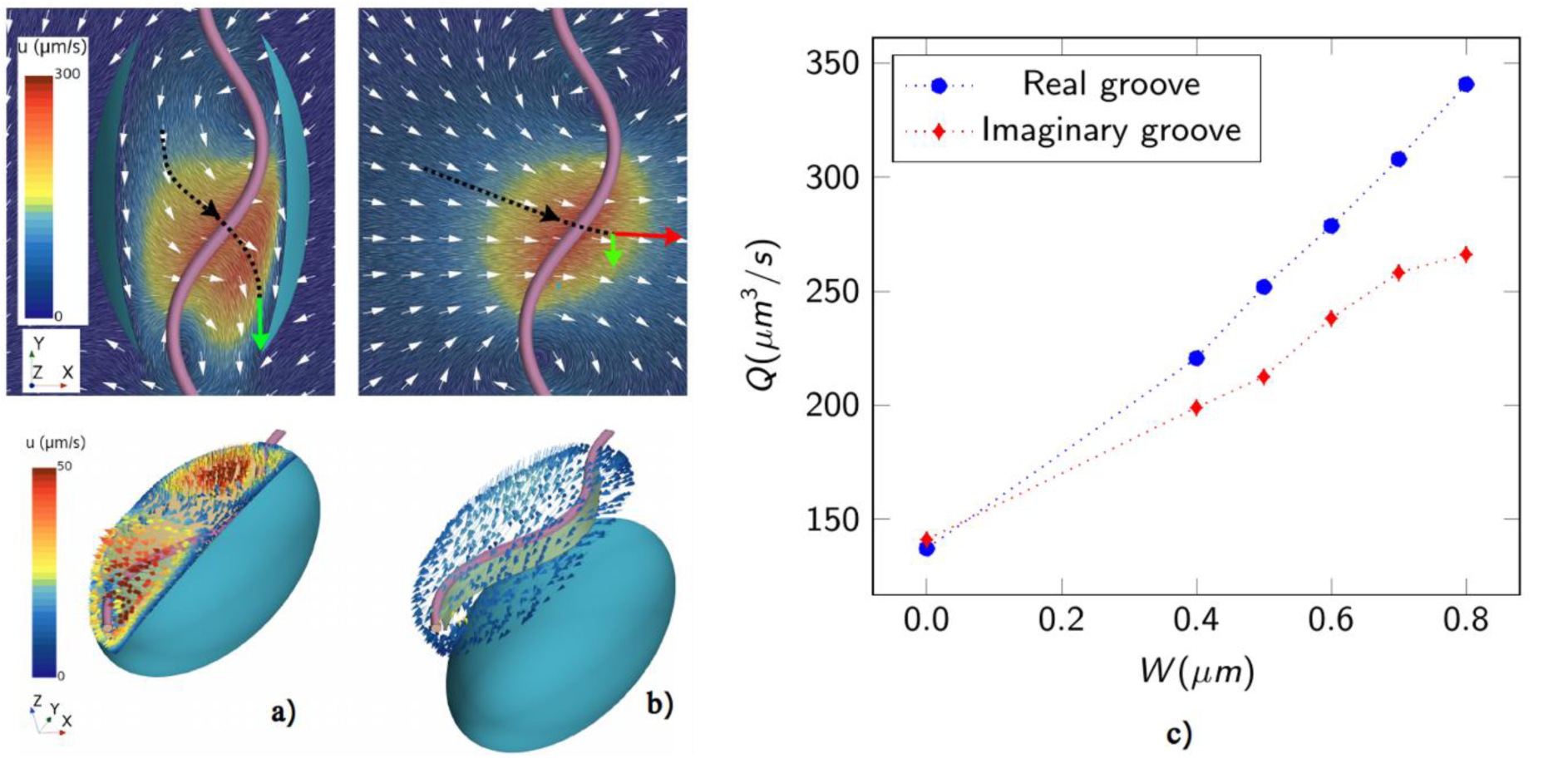
Effect of confinement on the feeding current generated by a (vaned) flagellum. A snapshot of the flow during the flagellum beating cycle (a, top), shows that the vaned flagellum pushes the flow against the wall of the groove, directing the flow posteriorly (green arrow). This interaction, during the complete beat cycle, results in a relatively strong averaged flow through the groove (a, bottom). In the absence of the groove (b, top), the vaned flagellum pushes the flow in free space where only a small component of such flow is directed downwards (green arrow), while most of the flow is directed sideways (red arrow). The oscillating sideways flows cancel out during the beat cycle, resulting in a weak average flow through an imaginary groove (b, bottom). The clearance rate (*Q*) increases with the width of the vane (*W*), but most so for a flagellum within a groove (c).

## Discussion

We have here described the suspension feeding and swimming behavior of several species of typical excavates drawn from across the three main phylogenetic groups. While phylogenetically diverse, they are all characterized by a vaned flagellum beating in a ventral groove. Obviously, this flagellar arrangement is primarily an adaptation to feeding, not swimming. In fact, like many other phagotrophic flagellates, excavates seem to be mainly attached to a surface when feeding, and free-swimming excavate cells appear to be slow and inefficient swimmers (Table 2). One exception among our study species is the swarmer stage of *R. americana* that adopts a different morphology and flagellar beat pattern than the attached foraging stage, and it lacks the flagellar vane (21). Also, the swimming form of the jakobid *Stygiella incarcerata* usually lacks a ventral groove and has a longer anterior flagellum (23). This contrasts with the less effective swimming behavior of the ‘normal grooved’ cells of *Jakoba libera* described here.

The clearance rates estimated here are of similar order of magnitude as previous estimates for *J. libera* derived from bottle incubation experiments] (24, 25), ∼250-800 µm^3^ s^-1^ (cf Table 4 and Fig. 5 for our experimental and computational estimates). These estimates, in turn, are within the typical range for phagotrophic flagellates of similar sizes (4).

Key to the functioning of the flagellum in clearing prey from ambient water is the interaction between the groove and the vaned flagellum. A vaned flagellum beating in a depression on the cell surface may be particularly efficient in generating a feeding current: First, having the force near the cell brings the feeding current streamlines near to the cell surface (Fig.s 4 and 5) where prey are captured, thus increasing prey encounter and retention rates. Second, adding a vane to a beating flagellum is an energy-efficient way of increasing the clearance rate, compared to increasing the beat frequency. The power consumption of a beating flagellum constrained in a groove increases linearly with its width, as does the resulting clearance rate, and the relation between power consumption and clearance rate is therefore near-linear (Fig. 7). In contrast, because in the viscous environment in which flagellates operate the clearance rate scales with the beat frequency while the power consumption with the beat frequency squared, the clearance rate increases only with the square root of the power consumption (Fig.7), rendering this a much less energy-efficient way of increasing the clearance rate. One may question the significance of the energy efficiency argument, because the operational cost of flagella typically constitutes only a tiny fraction of total metabolism of rapidly growing cells (26). However, flagellates may increase their activity but reduce their metabolism by orders of magnitude when resources are scarce (27) – a state in which they may be much of the time in nature – and in this situation, the energy efficiency of feeding may become important.

**Fig. 7.**
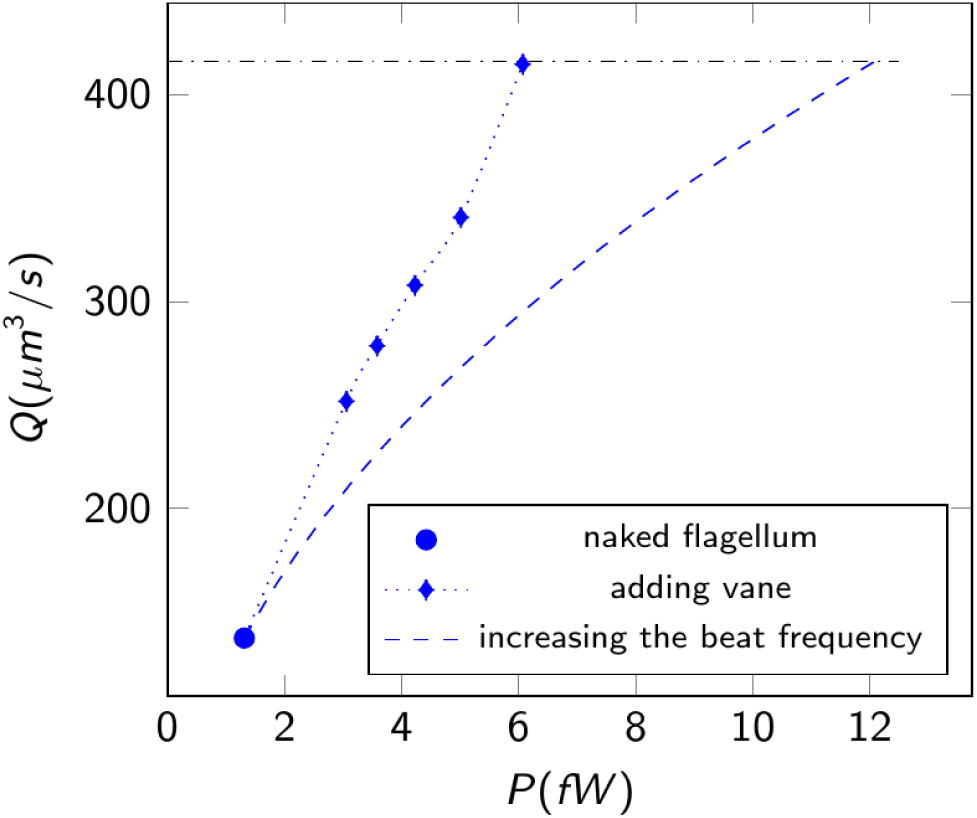
Clearance rate (Q) vs power consumption (P). Comparing clearance rates when adding vanes versus increasing the beat frequency. The data points represent simulation results for vane widths of 0.4µm, 0.5µm, 0.7µm, 0.8µm (1-vane configuration), and the last point 0.7µm (2-vane configuration). The dashed line illustrates the effect of increasing the beat frequency given by 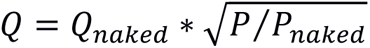.

Broadened (vaned) constrained flagella are widespread among phagotrophic flagellates. Choanoflagellates have a vaned flagellum beating inside a collar filter, and the equatorial flagellum of dinoflagellates drives a diaphragm-like ‘sock’ in the circumferential depression on the cell surface (cingulum), arrangements that both allow efficient feeding currents (5). These are not homologous structures, however. For example, the vanes in excavates are each supported by intraflagellar paraxonemal lamella, while the fibrillar vanes of choanoflagellates extend out of the flagellar membrane, and the cingulum-contained flagellum in dinoflagellates is equivalent to the anterior flagellum of excavates, rather than their posterior flagellum.

Conversely, there are other poorly studied phagotrophic flagellates from other deep branches of the eukaryotic tree of life with a flagellar arrangement similar and potentially homologous to that found in the typical excavates, namely a posterior flagellum with a vane supported by an intraflagellar lamellum that beats in association with a conspicuous ventral groove. The best examples are colponemids within the alveolates (28, 29) and the Nebulidia within the newest proposed supergroup – Provora (30, 31). The foraging mechanisms in these species are yet to be explored in detail, but the similarity of structures may suggest similar fluid dynamic behaviour, albeit both colponemids and nebulids are eukaryovores and ingest their prey at the anterior end of the cell, not the posterior (29).

The anterior flagellum and the ‘free’ part of the posterior flagellum apparently do not directly contribute to driving water through the groove. However, they may mix the ambient water and increase the spatial extension where the feeding current is effective, thus constantly providing fresh, prey-containing water to the immediate vicinity of the cell. This effect shares similarities with the mechanism observed in the pumping of choanocyte chambers in sponges (32). Such mixing may be relevant for cells attached to a surface where the viscous boundary layer may rapidly be depleted of prey. Typical excavates attached to a marine snow particle may not only be feeding on the elevated concentration of bacteria swarming around a particle (33), but may also be feeding on the high density of bacteria typically attached to the particle surface, as observed above for *C. membranifera* (34). Here again the interaction between the posterior flagellum and the groove appears important.

(a) Phylogenetic implications of functional morphology

In this study we examined species from all three principal clades of ‘excavates’ – Discoba, Metamonada and Malawimonadida. The foraging mechanisms appear fundamentally similar across the examined species, in terms of posterior flagellum activity, feeding current magnitude, and engulfment behavior. Arguably the greatest differences, between *Kipferlia* and the other subjects, divide species within a principal clade rather than between them, since *Kipferlia* and *Carpediemonas* are both metamonads. This functional similarity is broadly consonant with the hypothesis that the ‘typical excavate’ cell architecture is ultimately homologous across Discoba, Metamonada and Malawimonadida (16, 18, 35). The relationships among excavates are not well resolved, however, and most recent phylogenomic analyses show the three main clades representing at least two very distantly related lineages (16, 30). Reconciling the excavate homology hypothesis with these molecular phylogenetic results is possible if it is supposed that the Last Eukaryote Common Ancestor (LECA) was a ‘typical excavate’ in its morphology. Indeed, some form of excavate ancestry for all living eukaryotes has been advocated after some ‘rooted’ phylogenomic analyses of eukaryotes (19, 36). It is credible speculation, at least, that the typical excavate foraging system characterized here may have been used by our distant ancestors more than 1 billion years ago (37). This suggests an ancient history of bacterivory via relatively elaborate suspension feeding (rather than feeding solely by creeping on surfaces, for example), and permits reasonable estimates of the effectiveness of that feeding within ancient microbial ecosystems. Given these biological and earth ecosystem evolution stakes, more research into the details of typical excavate cell architecture, their extant diversity, and their phylogenetic relationships, is warranted.

## Authors’ contributions

All authors conceived the ideas, designed the study, interpreted the data, and contributed to the writing; SS, FM, and AGBS did the high-speed microscopic observations and interpreted the images; TK did the particle tracking; SSA developed the CFD model.

## Legends for online videos

**Movie S1.** *Jakoba libera* feeding mechanisms.

Movie S2. *Reclinomonas americana* feeding mechanisms.

Movie S3. *Kipferlia bialata* feeding mechanisms.

Movie S4. *Carpediemona membranifera* and *Malawimonas californiensis* feeding mechanisms.

Movie S5. Motion of vaned flagella in 5 species of excavate flagellates.

