## Supplementary material for "FORAGING MECHANISMS IN EXCAVATE FLAGELLATES SHED NEW LIGHT ON THE FUNCTIONAL ECOLOGY OF EARLY EUKARYOTES": online supplementary material

### SUPPLEMENTARY ONLINE MATERIAL

#### CFD Model Morphology:

Our generic computational fluid dynamics (CFD) model of excavates is a versatile model comprising a spheroid cell body with a depression (groove) on the cell body, and two flagella: a posterior flagellum and an anterior flagellum (figure S1). The cell has a minor axis length of 4.4µm, a major axis length of 7.8µm, and an initial orientation vector of [0.0, 1.0, 0.0], where the vector elements correspond to values along the X, Y, and Z axes, respectively. To create the groove, we extrude a cut in the shape of an ellipse positioned in a plane parallel to the XY-plane in the middle of the cell body. The ellipse has a minor axis length of 3.0µm, a major axis length of 7.8µm, and an orientation vector of [0.0, 1.0, 0.0]. The base of both the posterior and anterior flagella is positioned 2.2µm above the groove floor in the Z direction. The posterior flagellum has a planar beat, with the plane of the beat either parallel (base case) or perpendicular to the groove floor. In the base case, the length of the posterior flagellum is 9µm inside the groove but is also changed to 20µm by being extended (naked) outside the groove. A vane is positioned along 8µm of the flagellum (only inside the groove), starting from arc length 0.5µm from the flagellum base to avoid physical contact between the vane and the groove walls. The vane is always oriented perpendicular to the beat plane, either only on one side (1vane, base case) oriented inward (towards the groove floor) or outward, or on both sides (2vane). When present, the naked anterior flagellum has a length of 15µm and beats in a three-dimensional fashion. The beat frequency of both flagella is chosen f=35 Hz, and the kinematics of both the posterior and anterior flagella are provided in the supplementary materials. Finally, to model an excavate with a asymmetric groove walls, we rotate only the original spheroid such that its major axis has, for instance, the orientation of [0.1, 1.0, 0.0] as shown next.

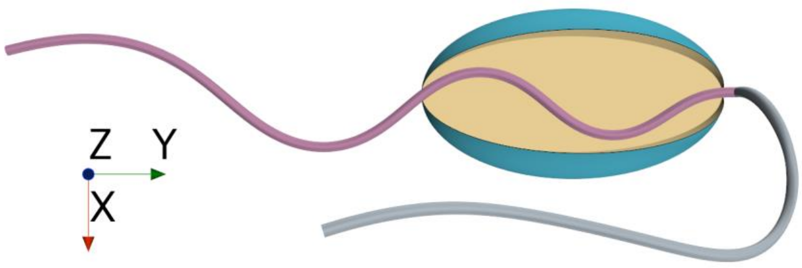


**Figure S1.** A versatile model depicting the morphology of excavates, featuring a spherical cell body (green) with a distinctive depression (yellow/gold), accompanied by a posterior flagellum (pink) and an anterior flagellum (silver). Various morphological parameters can be altered to encompass the diversity within excavates. These modifications encompass the removal of the anterior flagellum, exclusion of the "free" segment of the posterior flagellum (the portion outside the groove), addition of a vane on the posterior flagellum, adjustments to the beat plane orientation, the introduction of an asymmetric groove, and incorporating a surface in close proximity to the excavate.

**Symmetric and asymmetric groove:**

The versatile model morphology can accommodate both symmetric and asymmetric groove walls. In the symmetric case, the original spheroid has a major axis orientation of [0.00, 1.0, 0.00]. This configuration represents a groove where both sides of the groove walls are identical in shape and dimensions. However, in the asymmetric case, the major axis is oriented differently, such as [0.15, 1.0, 0.00], resulting in an uneven or asymmetrical groove as shown in figure S2.

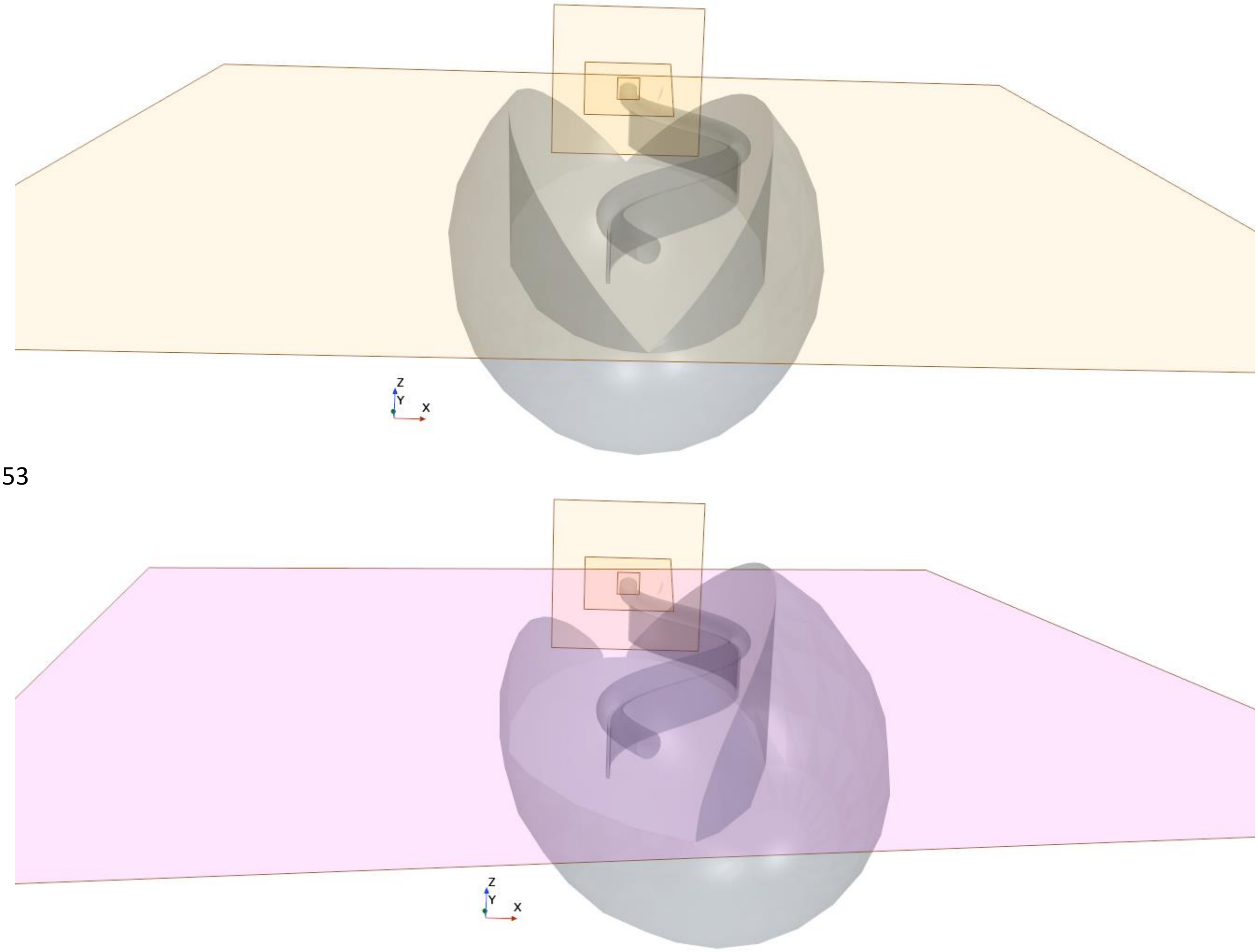


**Figure S2.** The modeling of symmetric and asymmetric groove walls. Top: The original spheroid with a major axis orientation of [0.00, 1.0, 0.00]. Bottom: The major axis oriented in [0.15, 1.0, 0.00].

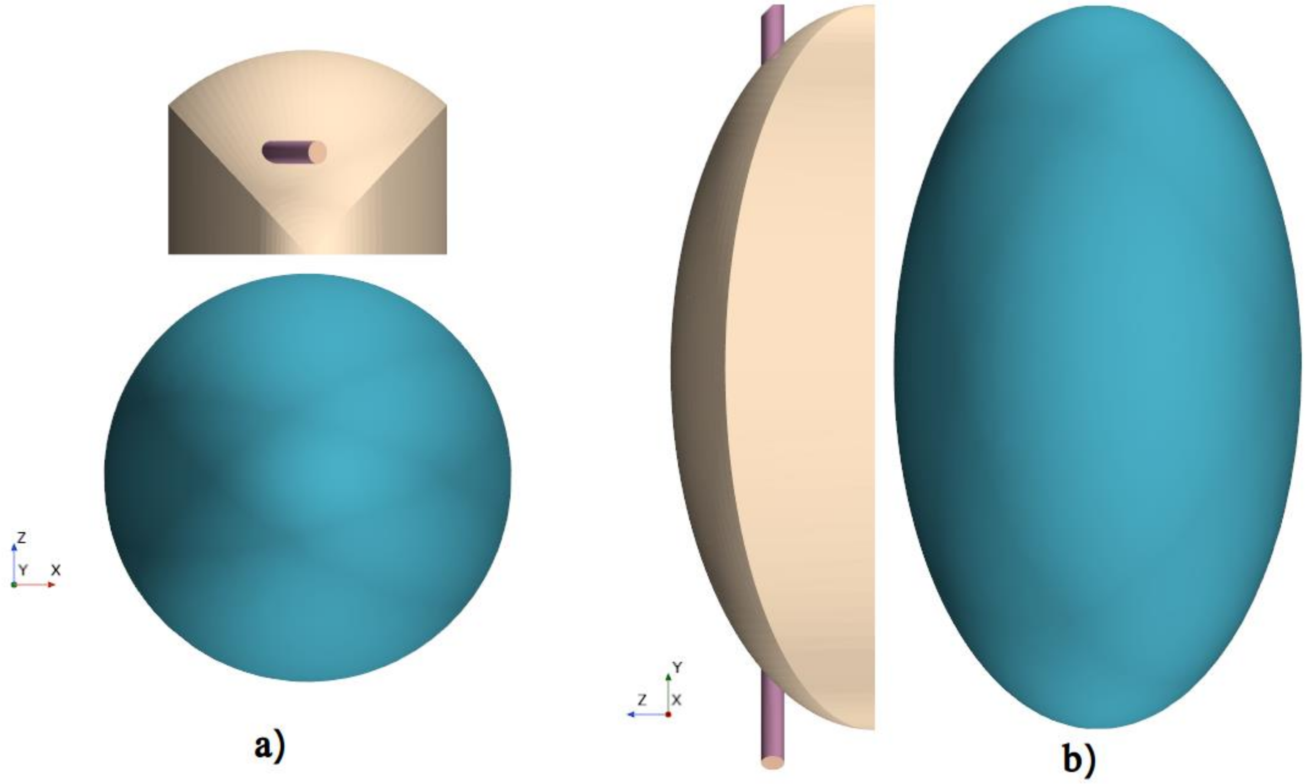


**Figure S3.** Two different views (a,b) depict the case with an imaginary groove (gold). The spheroid of the cell body (green) is displaced 2.3µm in the -Z direction, resulting in a groove-less spheroid cell. The shape of the imaginary groove mirrors the real groove in the base case before the displacement. The clearance rate into this imaginary groove is calculated for the 'imaginary groove' scenario.

**Computational Fluid Dynamics:**

We utilized the commercial CFD program STAR-CCM+ (version 18.02.008-R8) to numerically solve the Navier-Stokes equation and the equation of continuity for incompressible Newtonian flow. The finite-volume approach was employed for the simulation. The flow conditions were characterized by both a small frequency parameter and a low Reynolds number, resulting in a quasi-steady Stokes flow regime. Our computational model adopted the morphology of the organism, and a spherical domain with a diameter of 200µm was chosen which is ensured as large enough to not impact the results. The model cell was held stationary at the center of the domain. The no-slip boundary condition was applied to both the cell surfaces and the groove. The posterior and anterior flagella were modeled with a diameter of 0.25µm, while the vane had a thickness of 20 nm. All these components were subjected to the no-slip boundary condition. The flagellar beat was modeled as a planar wave for the posterior flagellum, and a 3-dimensional wave for the anterior flagellum. The waveform model for both posterior and anterior flagella in a general form is as follow:

$$\phi(s,t) = A_\phi\big(1 - exp(-s/\delta)\big)sin\big(2\pi ft - 2\pi\, s/\lambda_\phi\big) + CL_{fl}$$

where φ(s,t) represents the orientation angle of the flagellum tangent relative to the flagellar axis. The parameter 's' denotes the distance along the flagellum from its attachment point on the cell body, which is normalized by the length of the flagellum ($L_{fl}$). The variable 't' represents time. $A_\varphi$ corresponds to the amplitude of the angle, $\lambda_\varphi$ represents the wavelength, f indicates the beat frequency, and δ is an amplitude modulation factor that decreases the amplitude at the flagellum's attachment point to the cell body. C represents the turning angle of the flagellum, measured in radians (Geyer et al., 2016)

The beat pattern in the (x, y, z) domain presented here is derived from an expanded version of the planar waveform originally suggested by (Geyer et al., 2016). This extended waveform incorporates a more comprehensive three-dimensional pattern for the anterior flagellum. As a result, the (X(s), Y(s), Z(s)) coordinates of the flagellum are generated as follows:

$X(s) = \int_0^s dx,\ \ Y(s) = \int_0^s dY,$ and $Z(s) = \int_0^s dZ$

where

$$dx = \left(1/\sqrt{1 + C_z^2}\right)cos(\phi)ds,\ dy = \left(1/\sqrt{1 + C_z^2}\right)sin(\phi)ds,\ dz = \left(C_z/\sqrt{1 + C_z^2}\right)ds$$

which ensures that $ds^2 = dx^2 + dy^2 + dz^2$.

In the case of planar waveform of the posterior flagellum, the relevant parameters are, $A_\phi = 0.7, \delta = 0.1, \lambda_\phi = L_{fl}/3.2 + 2.0tan\big(\pi\, s/3\, L_{fl}\big), C_z = 0$

and in the case of 3-dimensional waveform of the anterior flagellum,

$$A_\phi = 1.25, \delta = 0.1, \lambda_\phi = 12.5, C_z = 5cos\left(\pi\big(s/L_{fl} + 0.3\big)\right).sin\left(0.5\pi\big(s/L_{fl}\big)\right)$$

We employ a combination of mesh morphing and the overset method to facilitate the movement of the computational mesh corresponding to the flagellar motion. The overset method deforms the mesh around the flagella, referred to as the overset region, rather than moving the entire mesh. This approach significantly reduces computational costs. Additionally, we generate a stationary background mesh that includes the cell, which overlaps with the deforming overset mesh. The two mesh regions are implicitly coupled, and field data are interpolated between them (i.e., the overset and background meshes) to generate a smooth solution at each iteration. By utilizing mesh morphing, we avoid the need to reconstruct the mesh geometry for different flagellum positions during the flagellar beat. The morphing motion redistributes mesh vertices in response to the flagellum's movement at each time step. As a result, the mesh undergoes morphing between time steps, aligning with the flagellar motion. At each time step, the discretized forms of the governing equations are solved within the entire computational domain. To accommodate the complex geometry of the model organism, we employ polyhedral cells for the discretization (figure S3), as they offer flexibility and enable mesh morphing.

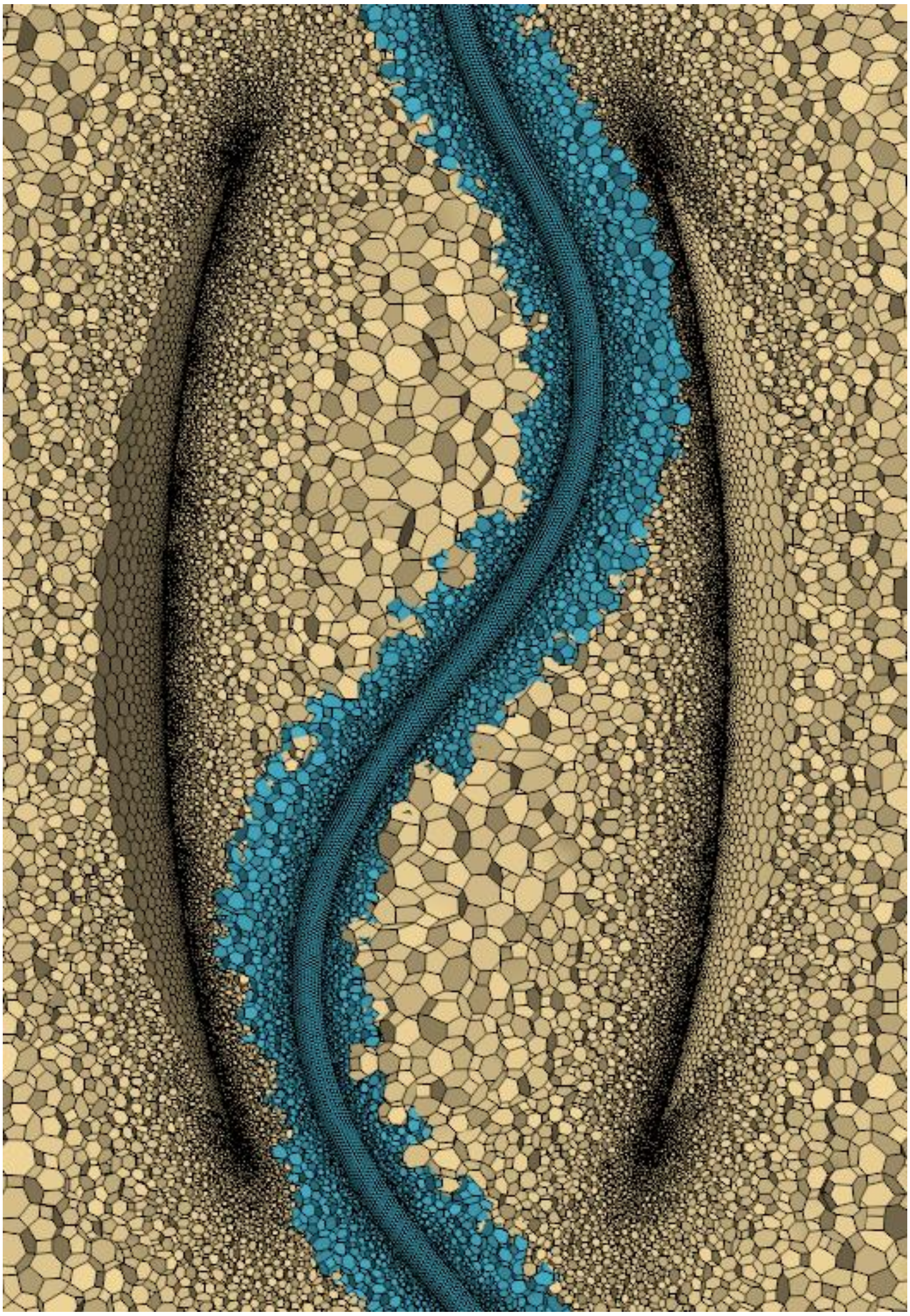

**Figure S4.** Visualization of the computational polyhedral cells in the beat plane. The computational domain consists of two separate regions, one including the (vaned) posterior and anterior flagella (overset mesh), and the other one stationary (background mesh) and includes the cell.
